# Protease mimicry: dissecting the ester bond crosslinking mechanics in bacterial adhesin proteins

**DOI:** 10.1101/2025.03.26.645365

**Authors:** Yuliana Yosaatmadja, Vanessa Ung, Xinlu Liu, Yixuan Zhao, Julia K. Wardega, Aria Shetty, Sophie Schoensee, Ivanhoe K. H. Leung, Jeremy R. Keown, David C. Goldstone, Edward N. Baker, Paul G. Young, Davide Mercadante, Christopher J. Squire

## Abstract

The ester bond crosslink discovered within bacterial adhesin proteins offers a captivating insight into the convergent evolution of enzyme-like machinery. Crystal structures reveal a putative catalytic triad comprising an acid-base-nucleophile combination and an oxyanion-like site that suggest a serine protease-like mechanism drives the crosslinking process. We now provide confirmation of the mechanism, revealing functional catalytic dyads or triads, and the recapitulation of protease machinery from a *Pseudomonas* bacterium and a human cytomegalovirus related only by convergent evolution. Molecular dynamics simulations show how a conservative threonine-to-serine mutation of the nucleophile induces hydrolysis and eliminates the ester bond crosslink. Collectively, our structural, functional, and computational efforts detail the molecular intricacies of intramolecular ester bond formation and underscore the convergent evolutionary adaptations of bacteria in exploiting enzyme-like machinery to protect essential adhesin proteins from the mechanical, biological, and chemical hostilities of the bacteria’s replicative niche.

## 1. Introduction

One of the first and the most critical steps in pathogenic bacterial infection is the adhesion of the pathogen to the host cell using long, filamentous surface appendages.^1–3^ Both Gram-negative and Gram-positive bacteria secrete surface adhesins that comprise single multidomain proteins or monomeric components for surface assembly of pili or fimbriae.^4–6^ Bacterial adhesins are key players in mediating host interaction and surface adhesion.^7^

In Gram-positive bacteria, “sticky” adhesin domains are located at the tip of hair-like pili or fimbriae that decorate the bacterial surface.^4; 8; 9^ Unlike the pili of Gram-negative bacteria that are multimeric protein assemblies held together by extensive non-covalent interactions,^10^ Gram-positive bacterial pili often consist of small Ig-like domains that are arranged like ‘beads on a string’ held together by intramolecular covalent bonds.^8^ These Gram-positive bacterial pili are narrow in diameter (20-30 Å) and are essentially one protein molecule wide when compared to their multimeric Gram-negative counterparts that measure 60-80 Å in diameter. ^4;9–12^ While these Gram-positive molecules are extremely thin, they can withstand persistent proteolytic, mechanical, and thermal stress during adhesion and host colonisation.^2; 13^ Central to this ability are non-canonical covalent intramolecular crosslinks between protein side chains forming isopeptide, thioester, and ester bonds that impart remarkable stability enhancement to single adhesin protein domains or are involved directly in covalent adhesion to host cells.^14–21^

While intramolecular isopeptide crosslinking in such domains requires only three amino acids buried in a hydrophobic environment, the ester bond equivalent has what appears to be a full complement of the catalytic machinery of a serine proteases enzyme, with up to six residues promoting nucleophilic attack between normally unreactive amino acid side chains.^14; 15; 22–27^ This configuration of enzyme-like features is conserved in the several X-ray crystal structures of homologous adhesin domains from Gram-positive bacteria, including human pathogens (*Clostridium perfringens,* PDB ID 4NI6 and 4MKM;^15^ *Mobiluncus mulieris,* PDB ID 5U5O and 5U6F),^28^ human oral commensal bacteria (*Gemella sp.,* PDB ID 7UC3 and 8F9L),^29^ and animal intestinal microbiota (*Enterococcus columbae* (pigeon), PDB ID 7UI8;^29^ *Suipraoptans intestinalis* (pig), PDB ID 8F90 and 8FHA).

The autocalytic machinery of ester bond crosslinking as first exemplified in the *Clostridium perfringens* adhesin Cpe0147, shows three residues (A577/H572/T450) arranged in space to mimic the acid/base/nucleophile catalytic triad of a serine protease, with Q580 the equivalent of a substrate electrophile (Figure 1).^15; 27^ Two buried and protonated acids, A480 and E547, act like the oxyanion site of a serine protease, polarising the carbonyl bond of Q580 side chain to increase its electrophilic potential and reactivity, and potentially stabilising a high energy tetrahedral intermediate. In the proposed mechanism, H572 abstracts a proton from T450 Oγ1, which then effects nucleophilic attack on the Q580 Cδ with the loss of the side chain amino group as ammonia – the final state is the equivalent of the acyl-intermediate of a serine protease mechanism, but without the potential to hydrolyse it is essentially trapped. The replacement of the nucleophile Thr-450 with a serine produces a crosslink species that is then hydrolysable at basic pH and we have further proposed that this reaction follows the second half of a serine protease mechanism whereby a water molecule attacks and breaks the ester bond of the acyl-intermediate.^30^

**Figure 1.**
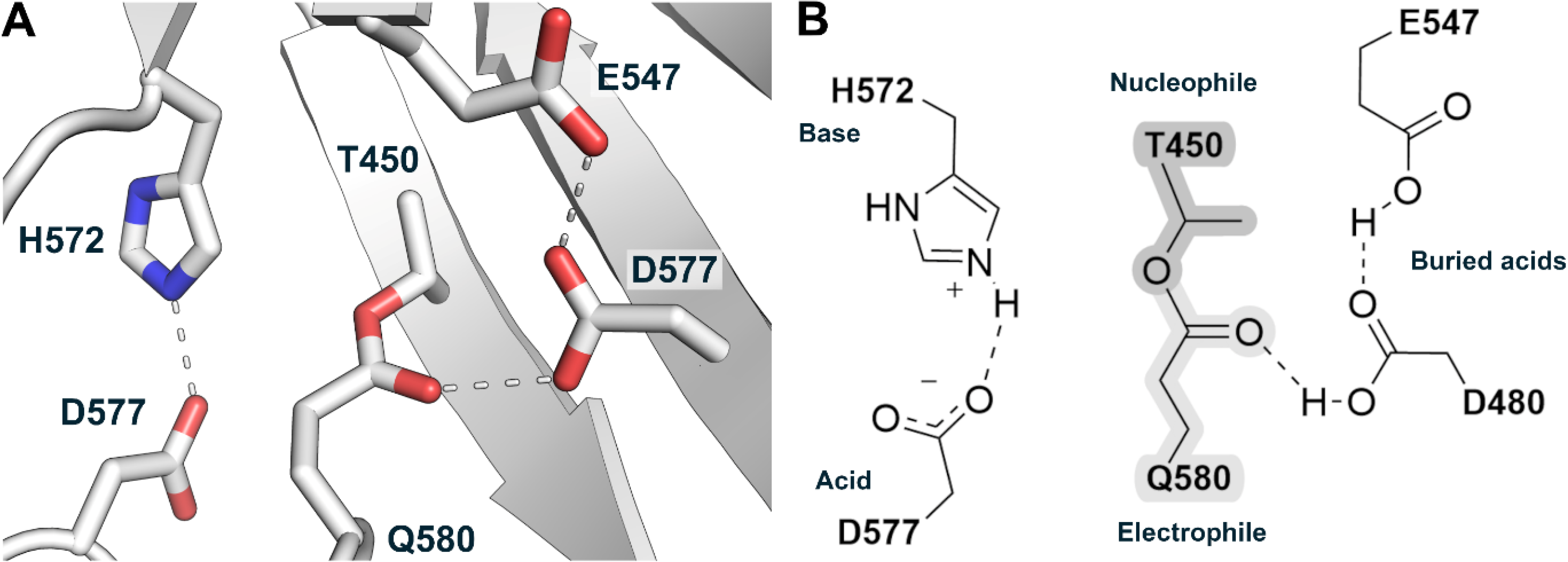
The autocatalytic site of Cpe0147. **A.** The arrangement of amino acid side chains around the intramolecular ester bond crosslink site in *C. perfringens* adhesin domain (PDB code 4NI6). The ester crosslink is formed in an autocatalytic reaction between a threonine and a glutamine that is proposed to follow a serine protease-like mechanism. **B.** 2D representation of the autocatalytic site with side chains labelled for their proposed functional role in catalysis as acid, base, nucleophile, or electrophile. The buried acids are equivalent to the oxyanion site of a serine protease and are proposed to stabilise a tetrahedral intermediate as well as the final crosslinked structure as shown. Dashed lines represent hydrogen bond interactions that may facilitate proton transfer during catalysis.

While this proposed ester bond-forming mechanism is compelling, is it a genuine example of convergent evolution? Using biochemical, biophysical, crystallographic, and computational studies, we can now reconcile previous experiments and validate a mechanism of ester bond formation in *Clostridium perfringens* Cpe0147 and homologous adhesin domains from Gram-positive bacteria. In the process of delineating the functional role of each amino acid of the proposed catalytic machinery, we have recapitulated the functional catalytic machinery of two atypical serine proteases, one from a *Pseudomonas* bacterium, the other a human herpes virus. The collective evidence strongly supports our proposition that the ester bond crosslinking machinery featured in bacterial adhesin domains is a *bona fide* example of convergent evolution of enzymatic mechanism tuned to bacterial virulence and survival.

## 2. Results

Our original site-directed mutagenesis studies described in Kwon *et al.* used a single Ig-like domain comprising the first repeat domain of the 12-domain *C. perfringens* adhesin protein Cpe0147.^15^ That study produced mutants without an ester bond crosslink that appeared unfolded by both differential scanning fluorimetry and circular dichroism. In the current study and following a closer examination of crystal structures (PDB ID 4NI6 and 4MKM), we designed a new construct comprising the second Ig-like domain (Cpe0147^439–587^) and including longer *N-* and *C-*terminal extensions to avoid potentially adverse domain truncation effects. This construct was used as the basis for a series of site-directed mutagenesis and electrophoresis, and X-ray crystallography experiments, and finally for molecular dynamics simulations to support the hypothesis of a serine protease-like mechanism of ester bond formation.

### 2.1 Changes to catalytic residues replicate the catalytic machinery of bacterial and viral serine proteases in nature

Mutations were made in wild-type Cpe0147^439–587^ protein to the proposed catalytic triad residues D557/H572/T450, the “substrate” Q580, and buried acids D480/E547, that correspond respectively to acid/base/nucleophile, electrophile, and oxyanion site features of a serine protease. All mutations tested compromise the catalytic efficiency but many changes produce viable catalytic machinery with activity between 5% and 95% of wild type (Figure 2).

**Figure 2.**
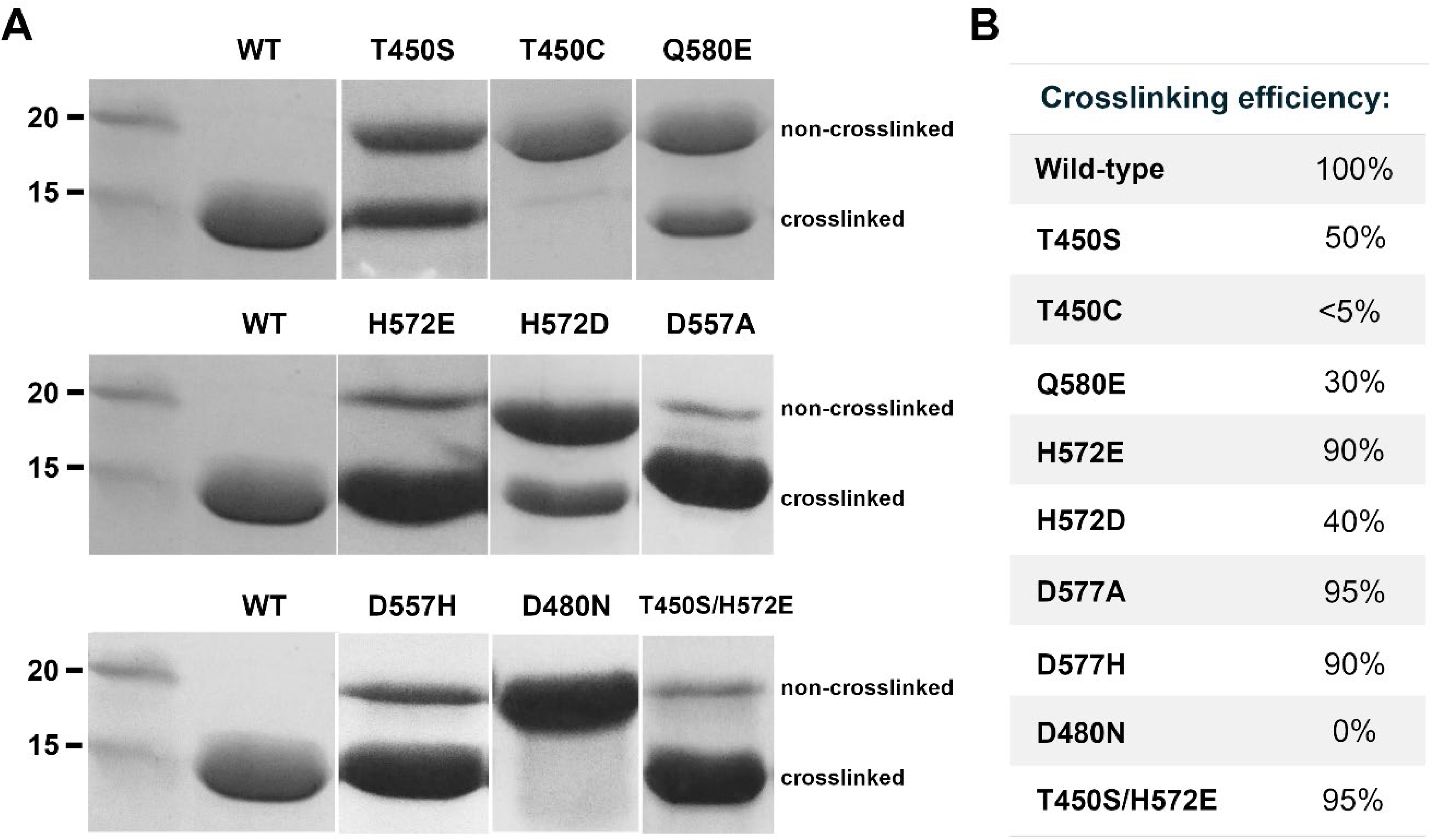
The effect of mutation on ester bond formation in Cpe0147. **A.** SDS-PAGE analysis of Cpe0147^439-587^ protein mutants relative to wild type (WT). Non-crosslinked protein migrates to ∼20 kDa and crosslinked protein to below 15 kDa apparent mass as indicated. **B.** Crosslinking efficiency – percentage of ester bond crosslink formed – as estimated by densitometry analysis of SDS-PAGE data using ImageJ.

In chymotrypsin and trypsin-like serine proteases, an aspartic acid is the first residue in the catalytic triad with multiple roles.^22; 31;32^ We proposed a similar set of roles for D577 in Cpe0147. To investigate this further, we produced alanine and histidine mutations of this aspartic acid (D577A and D577H). Both mutants produce an ester bond crosslink efficiently at between 90-95% of wild type (Figure 2). The D557A mutant suggests that a catalytic dyad is sufficient for catalysis. This is reiterated in the D557H protein that pairs two histidines as acid/base and replicates the His/His/Ser catalytic machinery of the human cytomegalovirus (CMV or human herpesvirus 5) protease where a dyad was also demonstrated as be sufficient for catalysis (Figure S1B).^33–36^

Mutation of the second putative triad residue H572 to alanine was previously shown to abolish ester bond formation and we suggested that this histidine was acting as both base and acid in catalysis.^15^ We now show that a glutamate can take the place of this histidine and retain close to wild-type cross-link efficiency (Figure 2; H572E, T450S/H572E). The T450S/H572E mutant reinforces the catalytic potential of a triad by replicating nature in the Asp/Glu/Ser catalytic triad of the serine protease sedolisin, found in *Pseudomonas* bacteria (Figure S1A).^37^ We searched the UniProt database^38^ for other occurrences of a glutamate as putative base in predicted ester bond adhesin domains. The search terms “VafE repeat-containing” and “T-Q ester bond” identified more than 1700 entries containing multiple ester bond repeat domains. Two entries for bacterial adhesins from *Agathobacter rectalis* and an unclassified *Lachnospiraceae* bacterium present an asparagine as the “acid” equivalent of D577 paired with a glutamate as base (Figure S2).

The nucleophile identity is sensitive to change with 100% crosslink formation in the wild-type threonine system dropping to 50% with a conservative serine substitution and further lowering to <5% for a cysteine substitution (Figure 2). T450S Cpe0147 has been characterised in detail previously as having a reversible crosslink that hydrolyses under specific buffer conditions. The mixture of species observed in Figure 2 may represent a dynamic equilibrium between crosslinking and hydrolysis.^30^ The T450C protein is found predominantly as non-crosslinked and disulfide crosslinked species under non-reducing SDS-PAGE (Figure 2) – this scenario is detailed below in the SEC-MALLS and X-ray crystallography results. The electrophile glutamine when mutated to glutamate (Q580E) and paired with wild-type nucleophile threonine, produces 30% crosslinked species (Figure 2).

Our final mutagenesis experiments probed the putative oxyanion site formed by the two buried acids, D480 and E547 that are proposed to stabilise a tetrahedral oxyanion intermediate via a series of hydrogen bonds or a proton shuttle arrangement. In previous mutagenesis studies, a D480A protein does not form the ester bond.^15^ Our D480N mutant also displayed a complete inability to form the ester crosslink (Figure 2). Taken together, these results suggest the absolute requirement of a buried protonated acid or oxyanion-like feature to effect autocatalysis in our crosslinking domains.

### 2.2 The dynamic behaviour of T450C in solution is linked to inefficient thioester crosslink formation

The hydrodynamic properties of wild type and T450C species in solution were investigated using SEC-MALLS. This technique has the advantage of separating mass calculation from shape or dynamic characteristics of the protein in solution. The wild-type protein displays a single species eluting at 12.5 mL with a calculated mass of ∼16 kDa as expected for a monomeric species (Figure 3A). By contrast, a more complex pattern of three elution peaks is produced by the T450C mutant with peaks at 9.7 mL, 11.0 mL, and 12.5 mL (Figure 3B). In agreement with SDS-PAGE, the largest species, eluting at 9.7 mL with a mass of 32 kDa, is a disulphide crosslinked dimer, followed by an non-crosslinked species eluting at 11 mL with a monomeric mass of 16 kDa (Figure 3B). In this system, by adding increasing concentrations of the reducing agent (TCEP), conversion of S-S crosslinked dimer to non-crosslinked monomeric protein is observed, again consistent with SDS-PAGE under reducing and non-reducing conditions. Only a very small proportion of T450C protein forms the smallest hydrodynamic radius, cross-linked species eluting similarly to the wild-type protein at ∼12.5 mL and with a thioester crosslink confirmed by mass spectrometry (Figure S3).

**Figure 3.**
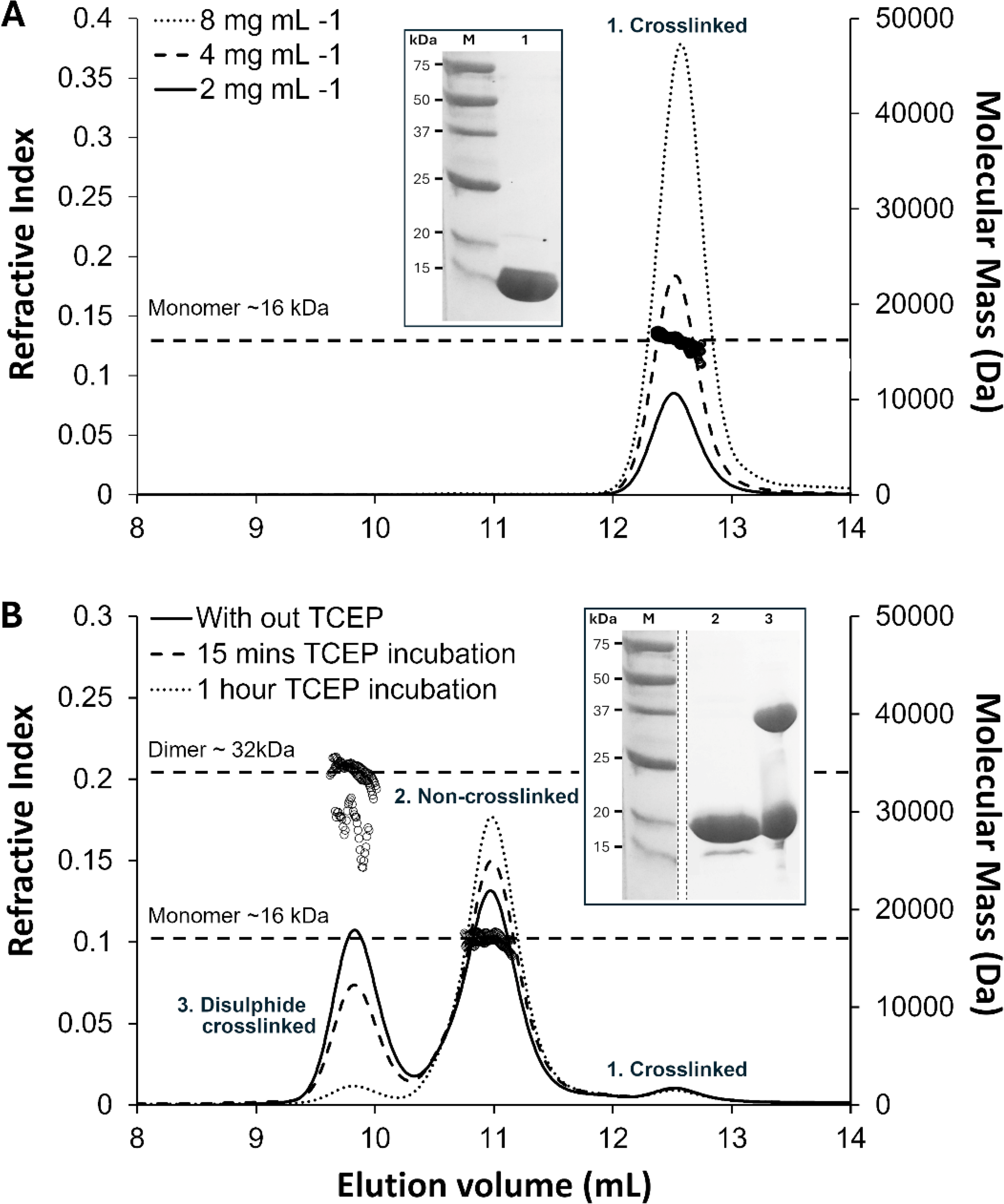
Characterisation of purified WT Cpe0147 and T450C mutant by SEC-MALLS and SDS-PAGE. **A.** SEC-MALLS trace of Cpe0147^439-587^ at three different concentrations shows a single crosslinked species eluting at 12.5 mL and with a diagnostic mass of 16 KDa. This is consistent with the reducing SDS-PAGE analysis (inset). **B**. SEC-MALLS trace of T450C mutant (5 mg mL^-1^) incubated with 2 mM TCEP for 0, 15, and 60 minutes as labelled. The trace displays three distinct peaks, from left-to-right, a disulphide crosslinked dimer (mass 32 KDa; elution vol. 9.8 mL), a non-crosslinked monomer (mass 16 KDa; elution vol. 11 mL), and crosslinked monomer (mass 16 KDa; elution vol. 12.5 mL). The result is consistent with SDS-PAGE analysis (inset) where a reducing environment shows 95% non-crosslinked and 5% crosslinked species, and a non-reducing has an additional disulphide crosslinked dimer with apparent mass of 37 KDa. Parts of the gel images have been removed for clarity.

### 2.3 The X-ray structure of the T450C mutant shows an unreactive nucleophile and provides a ground state model in a serine protease-like mechanism

The crystal structure of T450C Cpe0147^439–587^ (PDB 9BLO) was determined at 1.3 Å resolution with two molecules in the asymmetric unit (Supplementary Table 1). The two molecules are near-identical to the wild-type structure, with RMSD values of 0.53 Å and 0.44 Å, respectively, for Cα atoms of chain A and chain B of the mutant when compared to wild type (PDB 4NI6). A close inspection of the mutation site shows no thioester cross-link with the C450 thiol group sequestered in the pocket otherwise occupied by the methyl group of the wild-type T450 (Figure 4A). The reactive sidechain of Q580 forms hydrogen bonds through both Oε1 and Nε2 atoms with the protonated buried acid, D480. The position of E547 is unchanged compared to the wild type, pointing away from the ester bond position and making a hydrogen bond with D480.

**Figure 4.**
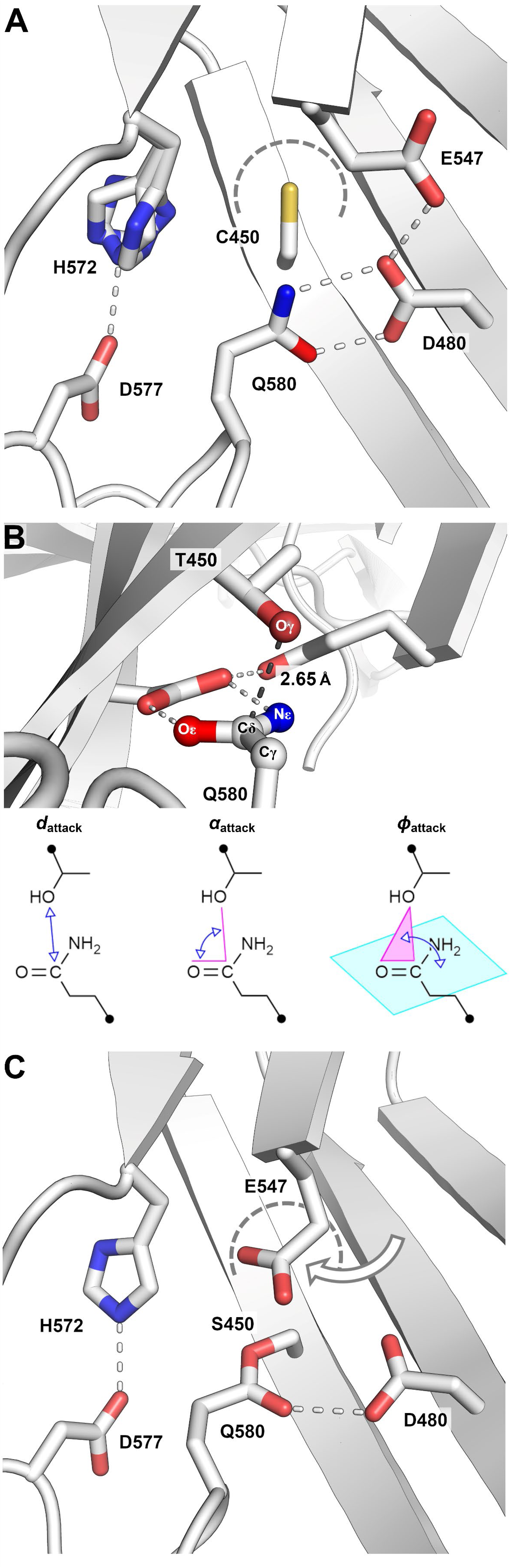
Crystal structures of Cpe0147^439-587^ T450C and T450S mutants support a serine protease-like mechanism of ester bond formation. **A.** The active site of the T450C mutant from chain B (PDB ID 9BLO) has no ester bond crosslink and the C450 thiol is rotated away from the bond-forming Q580 side chain into the pocket the WT T450 methyl group would occupy (dashed curved). Q580 forms a bidentate hydrogen bond to the buried acid D480. The histidine base, H572 adopts three conformations, one of which can hydrogen bond to its D577 partner. **B.** A ground state model of the WT crosslinking site modelled on the T450C structure. T450 Oγ closely approaches Q580 Cδ. The electrophile is held flat by hydrogen bonding to the buried acid and in a geometric arrangement consistent with that of serine proteases. Geometric parameters *d*_attack_, *α*_attack_, and *φ*_attack_ are defined in the same way as Du et al.^39^ **C.** The active site of the T450S mutant (PDB ID 9BLP)shows an ester bond crosslink and the buried E547 rotated towards the ester bond to fill space not occupied by a WT T450 methyl group (dashed curve). The E547 carboxylate/carboxylic acid functionality sits directly above the ester bond.

A wild-type, non-crosslinked model was produced from the T450C structure (Figure 4B). Geometric analysis following the method of Du *et al.* formulated for serine protease characterization, gives an attack distance, *d*_attack_, of 2.65 Å measured between nucleophile Oγ and electrophile target atom Cδ.^39^ The attack angles *α*_attack_ and *φ*_attack_ are calculated at 104° and 89°, respectively. Comparative average values for these parameters calculated from 1000+ serine protease crystal structures are *d*_attack_ = 2.68 (SD = 0.14 Å), *α*_attack_ = 93° (SD = 7°), and *φ*_attack_ = 84° (SD = 8°).

### 2.4 The T450S mutant X-ray structure shows a rearrangement of one of the proposed oxyanion-like buried acids

The crystal structure of T450S Cpe0147^439–587^ (PDB 9BLP) was determined at 1.2 Å (Supplementary Table 1) with one molecule in the asymmetric unit. The T450S protein, similarly to T450C, closely resembles the wild type with an RMSD comparison of 0.66 Å across all Cα atoms. An intramolecular ester bond linking S450 and Q580 is unambiguously present (Figure 4C). The pocket occupied by the T450 methyl group in wild type or thiol in T450C mutant, is partly occupied by the buried acid E547, which having rotated outwards from the wild-type localisation, no longer forms a proton shuttle with D480 and the ester bond carbonyl. Instead, E547 hydrogen bonds to an adjacent backbone carbonyl of V110 and stacks directly above the ester bond ether oxygen making a close contact (E547 Oε1 – S450 Oγ, 3.2 Å).

### 2.5 Molecular simulations explain the differences in hydrolysis seen for the S450 mutant and support a serine protease-like mechanism

We performed equilibrium molecular dynamics (MD) and metadynamics simulations of WT Cpe1047^439-587^ and a set of the experimentally tested mutants to further tease apart the role of putative catalytic triad residues and the susceptibility of the ester bond to hydrolysis when a serine nucleophile replaces threonine.

To recapitulate the proposed serine protease-like mechanism of Cpe0147^439-587^, we first simulated the D577H mutant to assess how D577 interacting with H572 would facilitate the correct positioning of H572 to act as a base for the proton abstraction of T450. In both WT and D577H proteins, the Nδ atom of H572 and the hydroxyl proton of T450 approach closely enough to effect protein abstraction (Figure S4A-B). Similarly, T450 is found in close proximity and in suitable geometries to form the ester bond with Q580. In both mutants, the dynamic behaviour of H572 relative to T450, and T450 relative to Q580, are near identical with the distance distributions predominantly overlapping in both systems (Figure S4C-D) and consistent with bond formation (Figure 2).

Additionally, we simulated the T450S/H572E double mutant, to assess if glutamate can act as acid/base in our mechanism. In line with the experimental bond formation efficiency (100% (WT) vs 95% crosslinked (T450S/H572E; Figure 2), the distribution of distances between E572 and S450, and between H572 and S450, overlap, with E572 even more closely approaching the nucleophile proton (Figure S5). A shift in the rotameric behaviour of E572 relative to the WT histidine (compare Figure S4H with S5H), shows the E572 sidechain often pointing towards the solvent, away from T/S450 (Figure S5G). This would overall suggest a reduced catalytic efficiency of T450S/H572E, as observed experimentally.

In a potential hydrolysis mechanism, the residue acting as acid to donate a proton back to the S450 nucleophile is unclear. The positioning of H572 in our crystals and comparison to a classic serine protease hydrolysis mechanism, suggests that this histidine could act as a catalytic acid to protonate the ester bond serine. However, our simulations suggest the key player in hydrolysis to be E547. In the WT structure, the carboxylic group of E547 is buried within the protein interior away from the ester bond, and is likely protonated and H-bonded to the second buried acid, D480. The T450S structure instead shows the E547 side chain rotated out of the protein interior and stacking directly above the ester bond ether’s oxygen (Ser-450 Oγ). By biasing E547 χ_1_ and χ_2_ dihedrals, metadynamics revealed the thermodynamics underlying the conformational space of E547, showing markedly different conformational energies between WT and T450S mutant. The free energy surface of WT Cpe1047^439-587^ features six low energy conformations, listed 1-6 in Figure 5B. Three conformations support hydrolysis (2, 4, and 5) with E547 stacked above the ester bond, and three (1, 3, and 6) have the side chain pointing away from the ester bond, unable to effect hydrolysis (Figure 5C). Well 6 has the lowest energy, preserving the WT H-bond with E480, and shows an energetic barrier of at least 28 kJ mol^-1^ relative to hydrolysis-promoting basins (Figure 5C). T→S substitution (Figure 5D-E) enhances the ability of E547 to swing towards the ester bond, to facilitate hydrolysis. In this case, E547 is unlikely to sample non-hydrolysing well 1, and the barrier between hydrolysing well 2 and non-hydrolysing well 3 is noticeably lowered (∼20 kJ mol^-1^) and unlikely hinders conformational switching. In T450S the most likely conformations are represented by hydrolysis-promoting basins 4 and 5 with E547 directly stacking above the ester bond. The differences between the threonine and serine-containing systems are evident in the distribution of distances between reactive groups in the different wells. For wells 4 and 5, the mean distance between E547 HE1 and S450 Oγ falls at approximately 3.5 Å, in line with what is observed in the crystal structure of this mutant. Conversely, the WT threonine offsets the same distance to higher values (Figure 5F).

**Figure 5:**
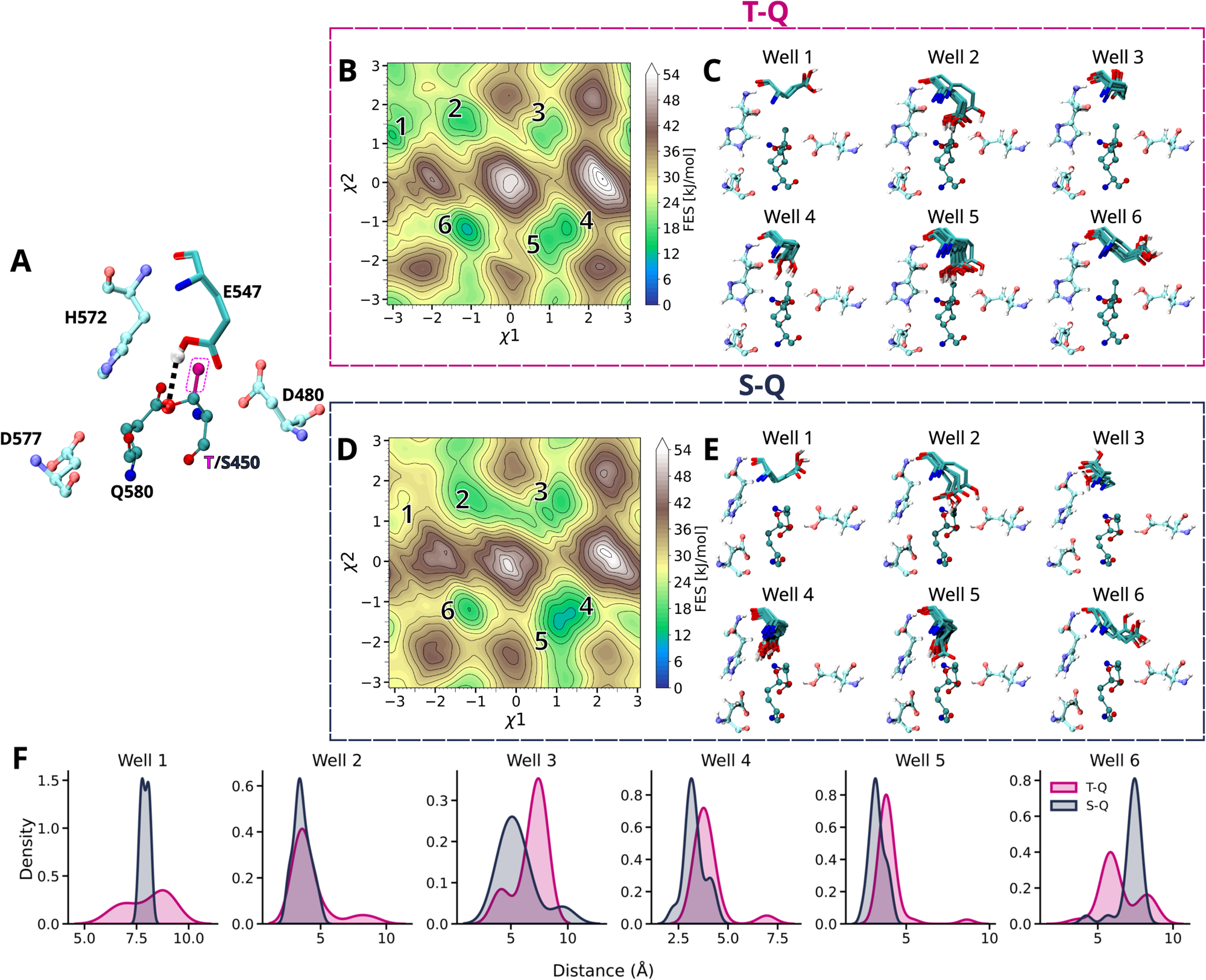
Simulations suggest why the T450S S-Q ester bond is more readily hydrolysed compared to a WT T-Q bond. **A.** Representative conformation of the T-Q covalent bonding and surrounding residues found in Cpe0147. E547 is highlighted for its ability to form a hydrogen bond with the ether oxygen of the ester bond in WT T450 protein (black dashed line). **B** and **D.** Two-dimensional free energy surfaces (FES) of the Thr-Gln or Ser-Gln simulated constructs, as obtained from metadynamics simulations using, as collective variables, the E547 χ1 and χ2 dihedral angles. **C** and **E.** Conformational ensembles of E547 isolated from the energy minima identified within the reconstructed FES (1 to 6 in panels **B** and **D**). **F**. Probability density distributions of the E547:HE1--T/S450:OG1 distance shown in **A** and obtained for each of the six identified energy minima.

## 3. Discussion

Adhesins in Gram-positive bacteria have evolved as single molecule wide constructions featuring intramolecular covalent crosslinks that have markedly changed our views on protein stability.^14; 15; 40^ We have proposed that the intramolecular ester bond crosslinking present in a subset of these adhesins represents a trapped acyl intermediate equivalent of a serine protease mechanism prior to hydrolysis.^41^ We have herein detailed structural, chemical, and dynamics factors determining ester bond formation in the *Clostridium perfringens* adhesin Cpe0147, and confirm a serine protease-like mechanism involving a catalytic triad (acid/base/nucleophile) or a dyad (base/nucleophile) and the equivalent of an oxyanion hole, as summarised in Figure 6.

**Figure 6.**
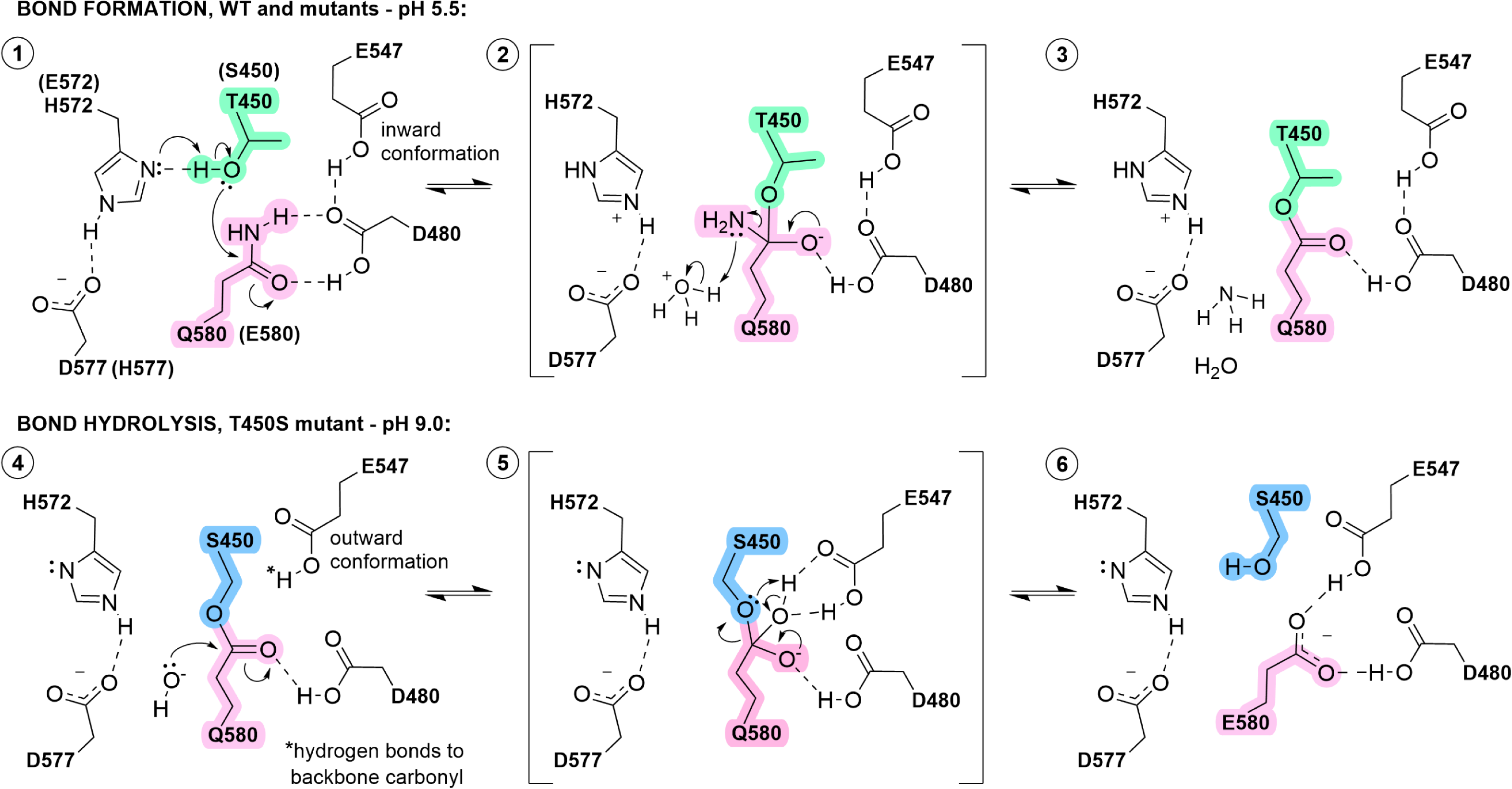
A refined mechanism of intramolecular ester bond formation and hydrolysis in bacterial adhesin domains. **Panel 1:** A wild-type model showing the position of the active site residues prior to ester bond formation was derived from the non-crosslinked T450C crystal structure (PDB 9BLO). One of the H572 rotamers derived from the T450C structure would be in hydrogen bonding distance to the WT T450 Oγ1 oxygen ready for proton abstraction. The bond forming E580 forms a bifurcated hydrogen bond to the buried acid residue D480. The Q580 Cδ atom is presented at an appropriate distance and geometry for nucleophilic attack by the T450 hydroxyl group. **Panels 2 and 3:** A proposed tetrahedral intermediate state formed under acidic conditions collapses to form the final crosslinked structure (PDB 4NI6) with the loss of ammonia. **Panel 4:** At pH 9, the S450 mutant undergoes attack by a hydroxide ion in solution. This configuration is modelled on the T450S crystal structure (PDB 9BLP) with E547 is an outward conformation. **Panels 5 and 6:** A tetrahedral intermediate is stabilised by both buried acids directly prior to its collapse. Proton abstraction from E547 regenerates the active site with E580 in place of the original Q580. Note that multiple steps, and electron and proton movements are combined and simplified; transition states are not shown. The revised mechanism is derived from that of Kwon *et al.* 2014 and is supported by X-ray crystal structures and MD simulations. Drawn using ChemDraw version 23.1.2 (Revvity Signals Software Inc.).

Perhaps the most compelling evidence for convergent evolution is the ground state configuration of WT protein derived from the T450C crystal structure (Figure 4B and Figure 6, panel 1). In this model, the close approach (2.65 Å) of nucleophile threonine Oγ and electrophile glutamine Cδ, and the perpendicular relative orientation of the side chains, is consistent with the geometric arrangement observed by Du et al. within 1000+ serine proteases.^39^ This close approach effectively destabilises the ground state of the system.

Du et al. further apportion the relative contribution of other elements towards the overall catalytic efficiency in serine proteases including the dominance of the catalytic base, contributing as much as half of the overall catalytic impetus, and the non-essential requirement of a catalytic acid.^39^ Similarly, in our system, the inclusion of either a histidine or glutamate (or aspartate to a lesser extent) as catalytic base appears essential. A catalytic dyad (minus the catalytic acid) appears equally viable given the D577A mutant retains 90% of the WT esterification. The D577H mutation that recapitulates the catalytic machinery of cytomegalovirus protease, removes the putative acid but retains a potential hydrogen bond donor atom. Studies by Khayat *et al*. suggested that the cytomegalovirus protease is likely to function via a catalytic dyad, but less efficiently compared to other serine proteases utilizing triads, and this correlates with our system.^42^

The recovery of T450S activity from 50% crosslinking efficiency to 95% by the addition of the second mutation, H572E, highlights the importance of a suitable base functionality in the putative catalytic dyad/triad. The T450S/H572E protein recapitulates the catalytic triad machinery of the serine protease sedolisin that is secreted by *Pseudomonas* species and which can hydrolyse insulin. Sedolisin is active between pH 3.0 and 5.0, bracketing the standard p*K*_a_ of a glutamic acid side chain of ∼ 4.1, allowing this side chain to function as acid or base as required by different steps in catalysis.^37; 43; 44^ While we did not test the pH dependence of our double mutant, our standard catalytic assay is completed at a relatively low pH of 5.5, close to the functional range of sedolisin. In search of a sedolisin-like glutamate base in nature, we identified more than 400 UniProt entries of putative bacterial adhesins containing predicted structures of ester bond crosslink domains. While crosslinks cannot be predicted by AlphaFold, the proximity and relative orientation of viable acid/base/nucleophile/electrophile and oxyanion-like hole residues, allowed us to confidently assign domains as homologous to the canonical Cpe0147 repeat domain and its catalytic machinery. Of note, a glutamate variation does appear within nature, however this was found in less than 0.5% of the UniProt entries surveyed. The two exemplars of the glutamate base in domains from *Agathobacter rectalis* and an unclassified *Lachnospiraceae* bacterium also display a paired asparagine as “acid” and further support the possibility of a functional catalytic dyad operating rather than a triad. The crosslinking potential of these putative adhesin domains remains to be validated experimentally.

In our experiments, the bond forming nucleophile/electrophile residues appear restricted to combinations of threonine/glutamine, threonine/glutamate, serine/glutamine, and serine/glutamate. While still catalytically viable, even conservative substitutions, for example, Q580E in the threonine/glutamate combination, greatly diminish bond formation, in this case to 50% of the wild-type yield. The differences we observe between the crosslinking ability of the two well-characterized nucleophiles cysteine and threonine, suggest a finely regulated crosslinking propensity, strongly dependent on the micro-environment of the reactive species. Cysteine is an obvious nucleophile involved in many biological reactions, including the cleavage of peptide bonds by cysteine proteases,^45^ the formation of thioester bonds in complement proteins,^46^ and the attachment of a raft of Gram-positive bacteria including *Clostridium perfringens* and *Streptococcus pyogenes* to host cells.^19; 21; 47^ However, a cysteine nucleophile in our system disfavours intramolecular crosslinking as the thiol group, already compromised by lower polarity than the WT hydroxyl, is also buried within the protein. A similar sequestration of the thiol group was noted in other serine protease-like enzymes where a catalytic threonine was replaced by a cysteine.^43; 44^ The highly dynamic solution properties of the non-crosslinked protein as indicated by SEC-MALLS and previously by NMR experiments,^30^ likely contributes to this lack of reactivity.

Our non-exhaustive survey of putative adhesin domains in the UniProt Database suggests that the vast majority of ester bond-forming sites in adhesin domains have evolved with a threonine as nucleophile, placing the methyl group in a pocket adjacent to the conformationally-restricted E547 and likely minimizing the potential for ester bond hydrolysis. Our simulations highlight a large energy barrier to E547 rotation. The methyl group of T450 closely interacts with the acyl chain of E547 and a large desolvation energy likely regulates the ability of E547 to rotate and attack the ester bond, as we see for the T450S system. However, in our adhesin domain-based molecular superglues from *Gemella bergeri* and *Enterococcus columbae* used extensively in our laboratory,^29^ a serine nucleophile does not promote hydrolysis even at pH 10 and when heated to 37-60°C (*unpublished work*). Additionally, a naturally occurring serine nucleophile-containing domain is predicted in the super-sized adhesin from *M. mulieris*, a bacterium naturally found in an acidic natural environment of pH 3-5.^28^ Assuming this native domain follows the pattern of the laboratory superglues in forming stable ester bond crosslinks, the autocatalytic site configuration in these systems is more complex than a simple substitution of a single amino acid in isolation, with the sites likely co-evolving in shape and chemistry to negate the hydrolysis potential.

Mechanistically, the ground state of bond formation emulates the active site configuration within the non-crosslinked C450 crystal structure and shows how the histidine or glutamate base deprotonates the T450 or S450 nucleophile (Figure 6, panel 1). MD simulations of H572 are characterized by subsets of well-defined conformers, some showing the Nδ of the histidine imidazolium ring approaching the T450 Oγ1 oxygen within distances and geometries optimal for abstraction. The role of the buried D480 as an oxyanion hole equivalent, is crucial for the bond formation, and following Du et al., we suggest its interactions with Q580 destabilise the ground state.^39^ Replacing this buried and presumably protonated aspartic acid with an asparagine, abolishes activity, both disrupting hydrogen bonding and removing a polarizing potential to the Q580 carbonyl bond. Stabilization of the putative oxyanion intermediate would also be ablated by this substitution. In the proposed intermediate state (Figure 6, panel 2), we suggest the involvement of a hydronium ion donating a proton to the leaving group, rather than the histidine that would play this role in a standard serine protease mechanism. This presumes that H572 is sequestered, as seen in the crystal structures of crosslinked WT and T450S domains, and is no longer in a suitable location and orientation for proton donation. Our experiments show that ester bond formation favours a pH <7.0 ^29; 30^ where hydronium ions are present in an excess, and this further supports our acid-mediated mechanism.

With respect to the mechanism of hydrolysis, as shown in the T450S crystal structure and reiterated by metadynamics simulations, the presence of the smaller serine relative to threonine considerably increases the likelihood of E547 directing its carboxyl group outward and close to the ether oxygen of the ester bond. At the higher pH values that promote hydrolysis, we propose that a hydroxide ion attacks the Q580 carbonyl to form an oxyanion intermediate stabilized by the outward facing, but still buried and presumably protonated E547 side chain (Figure 6, panels 4 and 5). The interactions formed with E547 would orientate the newly acquired hydroxyl functionality, facilitating the abstraction of the hydroxyl proton by the serine nucleophile as the tetrahedral intermediate collapses. The final state depicted in Figure 6, panel 6, represents a catalytic ground state of the serine protein ready for (re-)crosslinking if the pH is dropped to 5.5, but features a glutamate side chain in a hydrogen bond configuration different to that of the glutamine in panel 1. This altered and less perpendicular arrangement relative to the serine nucleophile, results in less efficient bond formation as observed for the Q590E mutant at 50% of WT efficiency (Figure 2).

Our findings reveal more of the intricate details behind ester bond formation within adhesin repeat domains. A catalytic triad remarkably similar in geometric configuration to serine proteases, most efficiently forms a crosslink. But our recreation of functional human cytomegalovirus protease-like machinery shows that a catalytic dyad can also drive esterification. Given the flexibility in catalytic residue choice seen across all serine proteases, it is unsurprising that more than one configuration of this system is also catalytically viable.^39;48; 49^ Like serine proteases, we see in crystal structures/simulations and propose in our refined catalytic mechanism, that coordinated motion of base, nucleophile, and electrophile promotes catalysis.^41^ These findings provide a compelling argument for convergent evolution in bacterial adhesin domains having “reimagined” serine protease machinery with a twist – by making bonds rather than breaking them.

## 4. Materials and methods

### 4.1 Cloning and site directed mutagenesis of WT and mutants of Cpe0147 (aa 439-587)

Constructs encoding Cpe0147^439–587^ (WT and mutants) were cloned into a modified pProExHta (Invitrogen) vector (pMBP-ProExHta), using EcoRI and KasI restriction endonucleases and T7 DNA ligase. pMBP-ProExHta was generated by inserting the *E. coli* maltose binding protein (MBP) gene between the His_6_-tag and the recombinant tobacco etch virus (TEV) protease (rTEV) cleavage site of pProExHta.^50^ Site-directed mutagenesis was carried out using a whole plasmid PCR method with complementary mutagenic primers,^51^ iProof enzyme (BioRad), DpnI treatment and T7 DNA ligase for re-circularisation before transformation into *E. coli* DH5α. Correct in-frame cloning and mutagenesis was confirmed by Sanger sequencing provided by the Auckland Genomics Centre, The University of Auckland, Auckland, New Zealand.

### 4.2 Expression and purification of WT and mutants of Cpe0147 (aa 439-587)

All Cpe0147^439-587^ WT and mutant constructs were transformed into *Escherichia coli BL*21 (DE3) cells and expressed at 18°C overnight in Terrific Broth medium supplemented with 50 mg mL^-1^ ampicillin and with shaking at 200 rpm. Cells were harvested by centrifugation, mechanically lysed, and subjected to immobilised metal affinity chromatography (IMAC), rTEV cleavage, and reverse IMAC treatments, as described previously.^30^ Proteins were concentrated to 1 mL in volume and purified by size-exclusion chromatography (SEC) using a Superdex 75 16/60 column equilibrated with SEC buffer (20 mM Tris.HCl pH 8.0, 100 mM NaCl and 2 mM βME). Fractions containing the target proteins were concentrated to 100 mg mL^-1^ and characterised by SDS-PAGE. The purified proteins were flash cooled in liquid nitrogen for later use.

The presence or absence of an intramolecular ester bond crosslink in the protein domains was assessed by SDS-PAGE with band intensity quantified using ImageJ^52^ and mass spectrometry – LC-MS/MS characterisation was performed as a service by the Mass Spectrometry Centre, The University of Auckland, Auckland, New Zealand.

### 4.3 Protein crystallization, data collection and refinement

The T450C Cpe0147^439-587^ was crystallised using sitting drop vapour diffusion using the Oryx 4 robot (Douglas Instruments) by mixing 200 nL protein with 200 nL of crystallization solution comprising 0.2 M MgCl_2_.6H_2_O, 0.1 M Tris.HCl pH 8.5, and 30% (w/v) PEG 4000. The T450S mutant Cpe0147^439-587^ was crystallised similarly from a solution comprising 0.2 M sodium thiocyanate and 20% PEG 3350. Both crystals were cryoprotected (crystallisation solution supplemented with 20% (v/v) glycerol), mounted in nylon loops, and flash cooled in liquid nitrogen.

X-ray diffraction data were collected at the Australian Synchrotron using the Blu-Ice software on the MX1 beamline, and were processed using XDS and AIMLESS ^25^.^53–55^ The structures were determined by molecular replacement in PHASER ^26^ using the structure of the wild-type protein (PDB coordinates 4NI6)^15; 56^ Iterative model building and refinement including the addition of water and ions used COOT and REFMAC.^57; 58^ Coordinates and structure factors were deposited in the Protein Data Bank^59^ with identifiers 9BLO and 9BLP for T450C and T450S structures respectively. Data collection and refinement statistics are given in Supplementary Table 1. A wild-type ground state model of the crosslinking site was produced from the T450C structure by mutating the cysteine to a threonine using COOT and then energy minimizing the coordinates. Geometric parameters *d*_attack_, *α*_attack_, and *φ*_attack_ were calculated from selected minimized atomic coordinates following the methodology of Du et al.^39^

### 4.4 Size-exclusion chromatography coupled to multi-angle laser light scattering (SEC-MALLS)

SEC-MALLS analysis was used to determine the molecular mass, cross-link presence/absence, and hydrodynamic properties of the proteins in solution. Protein (100 μL at 2-10 mg/mL) was loaded onto a Superdex 75 10/300 GL column equilibrated in 10 mM Tris.HCl pH 8.0, 150 mM NaCl, 3 mM NaN_3_, mounted on a Dionex HPLC with a PSS SLD7000 7-angle MALLS detector and Shodex RI-101 differential refractive index detector. The PSS winGPC Unicorn software determined averaged molecular masses for the eluted protein species.

### 4.5 Molecular dynamics and metadynamics simulations

Simulations were performed using Gromacs 2021.7. The protein was protonated as at pH 7.0 and placed in a box measuring 20 x 12 x 12 nm solvated with TIP3P water molecules. Na^+^ and Cl^-^ ions were added to achieve system neutrality to a total ionic strength of 0.15 M. Particles were assigned parameters from the Amber ff14SB force field.^60^ The system was firstly energy minimized to optimize atomic positioning, using a steep descent algorithm for 50000 steps with an energy step of 0.01 nm and a tolerance of 10 kJ mol^-1^ nm^-1^. Solvent equilibration was achieved in two steps: firstly, a 1 ns-long simulation was performed in the NVT ensemble with temperature kept constant at 300 K, coupled every 2.0 fs using a V-rescale thermostat.^61^ In this step, each atom was assigned random velocities, according to a Maxwell-Boltzmann distribution obtained at the simulated temperature. A second equilibration step was performed in the NPT ensemble keeping both temperature and pressure constant at 300 K and 1.0 bar, respectively. Pressure was coupled every 2.0 fs using the Parrinello-Rahman barostat.^62^ During both equilibration steps protein atoms were positionally restrained, applying a force constant of 1000 kJ mol^-1^ nm^-2^. A 500 ns-long run was then performed without positional restraints in the NPT ensemble.

Metadynamics simulations were performed using the last conformer of the equilibration run as the starting conformation. For each system (wild type and T450S mutant) three metadynamics simulation replicates were performed until convergence, for a total simulated time of 6.0 μs per run (Supplementary Table 2). The replicates differed in the set of initial velocities generated in the NVT equilibration step, as described above, and in the history of the bias added to explore the phase space along the chosen collective variables (χ_1_ and χ_2_ dihedral space). The rotameric space of the χ_1_ and χ_2_ dihedrals of residue 547 was explored and biased every ps by adding Gaussians of energy 0.2 kJ mol^-1^ high and 0.2 deg wide. The free energy profiles from the metadynamics simulations, were then obtained by reweighing each replicate to discount the bias accumulated during the simulations, according to the protocol described by Tiwary and Parrinello.^63^

A second set of molecular dynamics simulations, at equilibrium, were performed on D577H and T450S/H572E mutants, using a simulation protocol for box creation, solvation, and equilibration identical to that described above. Following equilibration, each system was simulated five times for 250 ns each (total simulated time 1.25 μs; Supplementary Table 2), with each replicate differing for the set of initial velocities assigned in the NVT equilibration step. The first 50 ns of each replica were discarded from analysis, as considered equilibration time. Protonation states were set to those occurring at pH 7.0, except for D480, E547, H572 and D577. D480 and E547 remained protonated for both mutants, while H572 and H577 in the D577H mutant were both singly protonated on the Nε atom. Residue E572 in the T450S/H572E mutant was left deprotonated.

All analyses were performed using a combination of *in-house* scripts on the collected trajectories and MDAnalysis.^64^ Representative structures were obtained using VMD 1.9.4.^65^

### 4.6 Bioinformatics and predicted structure analysis

A keyword search of the UniProt database (search terms; “T-Q ester bond”, “VafE repeat”) found more than 1500 protein entries. For those with AlphaFold predicted structures, ester bond crosslink domains and putative autocatalytic amino acids were identified by visual inspection using PyMol (Schrodinger) or Coot graphics software and by comparison to canonical ester bond domain Cpe0147 (PDB 4NI6).^57^

## Supplementary material description

Table 1. Data collection, refinement, and validation statistics for T450C and T450S X-ray crystal structures

Table 2. Molecular dynamics and metadynamics replicate and sampling parameters.

Figure 1. The active site triads of two non-cannonical serine proteases

Figure 2. Naturally occurring variations in the putative catalytic residues of Cpe0147-like ester crosslink domains.

Figure 3. Trypsin digest coupled with tandem mass spectrometry for T450C variant.

Figure S4: Stabilising interactions for T-Q bond formation in Cpe0147.

Figure S5: Stabilising interactions for S-Q bond formation in Cpe0147.

## Acknowledgements

This research was supported by the Marsden Fund Council from Government funding, managed by Royal Society Te Apārangi – grant UOA1421. This research was undertaken in part using the MX2 beamline at the Australian Synchrotron, part of ANSTO, and made use of the Australian Cancer Research Foundation (ACRF) detector. The authors wish to thank Martin Middleditch of the Mass Spectrometry Centre, The University of Auckland, Auckland, New Zealand for assistance with LC-MS/MS data acquisition and analysis. The authors wish to thank Kristen Boxen at the Auckland Genomics Centre, The University of Auckland, Auckland, New Zealand for assistance with Sanger sequencing.

## Description of supplementary material

File Squire-Protein-Science-supplementary.doc contains additional tables and figures. Table S1 – crystallographic data, Figure S1 – catalytic triad in two atypical serine proteases, Figure S2 – bioinformatics analysis of putative crosslink domains, Figure S3 – mass spectrometry analysis of thioester crosslinking, Figure S4 – molecular dynamics analysis of T-Q ester bond formation, Figure S5 – molecular dynamics analysis of T-S ester bond formation.

## Abbreviations and symbols

Ig-like: immunoglobulin-like
Cpe0147439-587: second repeat domain in surface-anchored adhesion protein from *Clostridium perfringens*
IMAC: immobilised metal affinity chromatography
SEC: size exclusion chromatography
SEC-MALLS: size exclusion chromatography with multi-angle laser light scattering
LC-MS/MS: liquid chromatography with tandem mass spectrometry
MBP: maltose binding protein
rTEV: recombinant tobacco etch virus protease.

## References

1. Pizarro-Cerda J, Cossart P. 2006. Bacterial adhesion and entry into host cells. Cell. 124(4):715–727.

2. Stones DH, Krachler AM. 2016. Against the tide: The role of bacterial adhesion in host colonization. Biochem Soc Trans. 44(6):1571–1580.

3. Krachler AM, Orth K. 2013. Targeting the bacteria-host interface: Strategies in anti-adhesion therapy. Virulence. 4(4):284–294.

4. Proft T, Baker EN. 2009. Pili in gram-negative and gram-positive bacteria - structure, assembly and their role in disease. Cell Mol Life Sci. 66(4):613–635.

5. Kline KA, Dodson KW, Caparon MG, Hultgren SJ. 2010. A tale of two pili: Assembly and function of pili in bacteria. Trends Microbiol. 18(5):224–232.

6. Sauer FG, Mulvey MA, Schilling JD, Martinez JJ, Hultgren SJ. 2000. Bacterial pili: Molecular mechanisms of pathogenesis. Curr Opin Microbiol. 3(1):65–72.

7. Baker EN, Squire CJ, Young PG. 2015. Self-generated covalent cross-links in the cell-surface adhesins of gram-positive bacteria. Biochem Soc Trans. 43(5):787–794.

8. Ton-That H, Schneewind O. 2004. Assembly of pili in gram-positive bacteria. Trends Microbiol. 12(5):228–234.

9. Kang HJ, Baker EN. 2012. Structure and assembly of gram-positive bacterial pili: Unique covalent polymers. Curr Opin Struct Biol. 22(2):200–207.

10. Walczak MJ, Puorger C, Glockshuber R, Wider G. 2014. Intramolecular donor strand complementation in the e. Coli type 1 pilus subunit fima explains the existence of fima monomers as off-pathway products of pilus assembly that inhibit host cell apoptosis. J Mol Biol. 426(3):542-549.

11. Craig L, Volkmann N, Arvai AS, Pique ME, Yeager M, Egelman EH, Tainer JA. 2006. Type iv pilus structure by cryo-electron microscopy and crystallography: Implications for pilus assembly and functions. Mol Cell. 23(5):651–662.

12. Waksman G, Hultgren SJ. 2009. Structural biology of the chaperone-usher pathway of pilus biogenesis. Nat Rev Microbiol. 7(11):765–774.

13. Dufrene YF, Viljoen A. 2020. Binding strength of gram-positive bacterial adhesins. Front Microbiol. 11:1457.

14. Kang HJ, Coulibaly F, Clow F, Proft T, Baker EN. 2007. Stabilizing isopeptide bonds revealed in gram-positive bacterial pilus structure. Science. 318(5856):1625–1628.

15. Kwon H, Squire CJ, Young PG, Baker EN. 2014. Autocatalytically generated thr-gln ester bond cross-links stabilize the repetitive ig-domain shaft of a bacterial cell surface adhesin. Proc Natl Acad Sci U S A. 111(4):1367–1372.

16. Kang HJ, Baker EN. 2009. Intramolecular isopeptide bonds give thermodynamic and proteolytic stability to the major pilin protein of streptococcus pyogenes. J Biol Chem. 284(31):20729–20737.

17. Alegre-Cebollada J, Badilla CL, Fernandez JM. 2010. Isopeptide bonds block the mechanical extension of pili in pathogenic streptococcus pyogenes. J Biol Chem. 285(15):11235–11242.

18. Pointon JA, Smith WD, Saalbach G, Crow A, Kehoe MA, Banfield MJ. 2010. A highly unusual thioester bond in a pilus adhesin is required for efficient host cell interaction. J Biol Chem. 285(44):33858–33866.

19. Walden M, Edwards JM, Dziewulska AM, Bergmann R, Saalbach G, Kan SY, Miller OK, Weckener M, Jackson RJ, Shirran SL et al. 2015. An internal thioester in a pathogen surface protein mediates covalent host binding. Elife. 4.

20. Lei H, Ma Q, Li W, Wen J, Ma H, Qin M, Wang W, Cao Y. 2021. An ester bond underlies the mechanical strength of a pathogen surface protein. Nat Commun. 12(1):5082.

21. Miller OK, Banfield MJ, Schwarz-Linek U. 2018. A new structural class of bacterial thioester domains reveals a slipknot topology. Protein Sci. 27(9):1651–1660.

22. Blow DM, Birktoft JJ, Hartley BS. 1969. Role of a buried acid group in the mechanism of action of chymotrypsin. Nature. 221(5178):337–340.

23. Dodson G, Wlodawer A. 1998. Catalytic triads and their relatives. Trends Biochem Sci. 23(9):347–352.

24. Polgar L, Bender ML. 1969. The nature of general base-general acid catalysis in serine proteases. Proc Natl Acad Sci U S A. 64(4):1335–1342.

25. Carter P, Wells JA. 1988. Dissecting the catalytic triad of a serine protease. Nature. 332(6164):564–568.

26. Hedstrom L. 2002. Serine protease mechanism and specificity. Chemical Reviews. 102(12):4501–4524.

27. Schwarz-Linek U, Banfield MJ. 2014. Yet more intramolecular cross-links in gram-positive surface proteins. Proc Natl Acad Sci U S A. 111(4):1229–1230.

28. Young PG, Paynter JM, Wardega JK, Middleditch MJ, Payne LS, Baker EN, Squire CJ. 2023. Domain structure and cross-linking in a giant adhesin from the mobiluncus mulieris bacterium. Acta Crystallogr D Struct Biol. 79(Pt 11):971–979.

29. Young PG, Squire CJ. 2020. Molecular superglues: Discovery and engineering orthogonalization. Methods Mol Biol. 2073:85–99.

30. Young PG, Yosaatmadja Y, Harris PW, Leung IK, Baker EN, Squire CJ. 2017. Harnessing ester bond chemistry for protein ligation. Chem Commun (Camb). 53(9):1502–1505.

31. Craik CS, Roczniak S, Largman C, Rutter WJ. 1987. The catalytic role of the active site aspartic acid in serine proteases. Science. 237(4817):909–913.

32. Kasserra HP, Laidler KJ. 1969. Mechanisms of action of trypsin and chymotrypsin. 47(21):4031–4039.

33. Chen P, Tsuge H, Almassy RJ, Gribskov CL, Katoh S, Vanderpool DL, Margosiak SA, Pinko C, Matthews DA, Kan CC. 1996. Structure of the human cytomegalovirus protease catalytic domain reveals a novel serine protease fold and catalytic triad. Cell. 86(5):835–843.

34. Qiu X, Culp JS, DiLella AG, Hellmig B, Hoog SS, Janson CA, Smith WW, Abdel-Meguid SS. 1996. Unique fold and active site in cytomegalovirus protease. Nature. 383(6597):275–279.

35. Shieh HS, Kurumbail RG, Stevens AM, Stegeman RA, Sturman EJ, Pak JY, Wittwer AJ, Palmier MO, Wiegand RC, Holwerda BC et al. 1996. Three-dimensional structure of human cytomegalovirus protease. Nature. 383(6597):279–282.

36. Tong L, Qian C, Massariol MJ, Bonneau PR, Cordingley MG, Lagace L. 1996. A new serine-protease fold revealed by the crystal structure of human cytomegalovirus protease. Nature. 383(6597):272–275.

37. Wlodawer A, Li M, Dauter Z, Gustchina A, Uchida K, Oyama H, Dunn BM, Oda K. 2001. Carboxyl proteinase from pseudomonas defines a novel family of subtilisin-like enzymes. Nat Struct Biol. 8(5):442–446.

38. Consortium TU. 2024. Uniprot: The universal protein knowledgebase in 2025. Nucleic Acids Research. 53(D1):D609–D617.

39. Du S, Kretsch RC, Parres-Gold J, Pieri E, Cruzeiro VWD, Zhu M, Pinney MM, Yabukarski F, Schwans JP, Martínez TJ et al. Conformational ensembles reveal the origins of serine protease catalysis. Science. 387(6735):eado5068.

40. Kang HJ, Paterson NG, Gaspar AH, Ton-That H, Baker EN. 2009. The corynebacterium diphtheriae shaft pilin spaa is built of tandem ig-like modules with stabilizing isopeptide and disulfide bonds. Proc Natl Acad Sci U S A. 106(40):16967–16971.

41. Radisky ES, Lee JM, Lu CJ, Koshland DE, Jr. 2006. Insights into the serine protease mechanism from atomic resolution structures of trypsin reaction intermediates. Proc Natl Acad Sci U S A. 103(18):6835–6840.

42. Khayat R, Batra R, Massariol MJ, Lagace L, Tong L. 2001. Investigating the role of histidine 157 in the catalytic activity of human cytomegalovirus protease. Biochemistry. 40(21):6344–6351.

43. Wlodawer A, Li M, Gustchina A, Oyama H, Dunn BM, Oda K. 2003. Structural and enzymatic properties of the sedolisin family of serine-carboxyl peptidases. Acta Biochim Pol. 50(1):81–102.

44. Reichard U, Lechenne B, Asif AR, Streit F, Grouzmann E, Jousson O, Monod M. 2006. Sedolisins, a new class of secreted proteases from aspergillus fumigatus with endoprotease or tripeptidyl-peptidase activity at acidic phs. Appl Environ Microbiol. 72(3):1739–1748.

45. Otto HH, Schirmeister T. 1997. Cysteine proteases and their inhibitors. Chem Rev. 97(1):133–172.

46. Pangburn MK. 1992. Spontaneous thioester bond formation in alpha 2-macroglobulin, c3 and c4. FEBS Lett. 308(3):280-282.

47. Linke-Winnebeck C, Paterson NG, Young PG, Middleditch MJ, Greenwood DR, Witte G, Baker EN. 2014. Structural model for covalent adhesion of the streptococcus pyogenes pilus through a thioester bond. J Biol Chem. 289(1):177–189.

48. Ekici OD, Paetzel M, Dalbey RE. 2008. Unconventional serine proteases: Variations on the catalytic ser/his/asp triad configuration. Protein Sci. 17(12):2023–2037.

49. Polgar L. 2005. The catalytic triad of serine peptidases. Cell Mol Life Sci. 62(19-20):2161–2172.

50. Ting YT, Batot G, Baker EN, Young PG. 2015. Expression, purification and crystallization of a membrane-associated, catalytically active type i signal peptidase from staphylococcus aureus. Acta Crystallogr F Struct Biol Commun. 71(Pt 1):61–65.

51. Dominy CN, Andrews DW. 2003. Site-directed mutagenesis by inverse pcr. Methods Mol Biol. 235:209–223.

52. Schneider CA, Rasband WS, Eliceiri KW. 2012. Nih image to imagej: 25 years of image analysis. Nature Methods. 9(7):671–675.

53. Kabsch W. 2010. Xds. Acta Crystallogr D Biol Crystallogr. 66(Pt 2):125–132.

54. McPhillips TM, McPhillips SE, Chiu HJ, Cohen AE, Deacon AM, Ellis PJ, Garman E, Gonzalez A, Sauter NK, Phizackerley RP et al. 2002. Blu-ice and the distributed control system: Software for data acquisition and instrument control at macromolecular crystallography beamlines. J Synchrotron Radiat. 9(Pt 6):401–406.

55. Evans PR, Murshudov GN. 2013. How good are my data and what is the resolution? Acta Crystallogr D Biol Crystallogr. 69(Pt 7):1204–1214.

56. McCoy AJ, Grosse-Kunstleve RW, Adams PD, Winn MD, Storoni LC, Read RJ. 2007. Phaser crystallographic software. J Appl Crystallogr. 40(Pt 4):658–674.

57. Emsley P, Cowtan K. 2004. Coot: Model-building tools for molecular graphics. Acta Crystallogr D Biol Crystallogr. 60(Pt 12 Pt 1):2126-2132.

58. Murshudov GN, Skubak P, Lebedev AA, Pannu NS, Steiner RA, Nicholls RA, Winn MD, Long F, Vagin AA. 2011. Refmac5 for the refinement of macromolecular crystal structures. Acta Crystallogr D Biol Crystallogr. 67(Pt 4):355–367.

59. Berman H, Henrick K, Nakamura H. 2003. Announcing the worldwide protein data bank. Nat Struct Biol. 10(12):980.

60. Maier JA, Martinez C, Kasavajhala K, Wickstrom L, Hauser KE, Simmerling C. 2015. Ff14sb: Improving the accuracy of protein side chain and backbone parameters from ff99sb. J Chem Theory Comput. 11(8):3696–3713.

61. Berendsen HJC, Postma JPM, van Gunsteren WF, DiNola A, Haak JR. 1984. Molecular dynamics with coupling to an external bath. The Journal of Chemical Physics. 81(8):3684–3690.

62. Parrinello M, Rahman A. 1981. Polymorphic transitions in single crystals: A new molecular dynamics method. Journal of Applied Physics. 52(12):7182–7190.

63. Tiwary P, Parrinello M. 2015. A time-independent free energy estimator for metadynamics. The Journal of Physical Chemistry B. 119(3):736–742.

64. Gowers RJ, Linke M, Barnoud J, Reddy TJE, Melo MN, Seyler SL, Domanski J, Dotson DL, Buchoux S, Kenney IM et al. 2019. Mdanalysis: A python package for the rapid analysis of molecular dynamics simulations. Paper presented at: Conference: PROC OF THE 15th PYTHON IN SCIENCE CONF (SCIPY 2016) ; 2016-07-11 - 2016-07-11 ;. Los Alamos National Laboratory (LANL), Los Alamos, NM (United States); United States.

65. Michaud-Agrawal N, Denning EJ, Woolf TB, Beckstein O. 2011. Mdanalysis: A toolkit for the analysis of molecular dynamics simulations. 32(10):2319–2327.

