## Supplementary Material for "Protease mimicry: dissecting the ester bond crosslinking mechanics in bacterial adhesin proteins"

| | T450C | T450S |
|---|---|---|
| PDB identifier | 9BLO | 9BLP |
| **Data Collection and processing:** | | |
| Diffraction source | MX1, Australian Synchrotron | MX1, Australian Synchrotron |
| Wavelength (Å) | 0.9537 | 0.9537 |
| Space group | *P*1 | *C*2 |
| Cell dimensions: | | |
| *a, b, c* (Å) | 39.98, 44.40, 50.1 | 54.91 44.17 64.70 |
| *α, β, γ* (°) | 95.1, 109.4, 110.1 | 90.00 110.1 90.00 |
| Resolution (Å)* | 18.86 – 1.35 (1.37 – 1.35) | 17.87 - 1.20 (1.22 - 1.20) |
| CC1/2* | 1.000 (0.79) | 0.999 (0.95) |
| $I/\sigma(I)$* | 27.8 (2.4) | 20.2 (3.8) |
| Completeness (%)* | 94.8 (77.4) | 97.7 (93.9) |
| Multiplicity* | 3.3 (3.3) | 7.1 (6.8) |
| **Refinement:** | | |
| Resolution (Å) | 18.86 – 1.35 | 17.87 - 1.20 |
| No. of reflections | 58614 | 42278 |
| $R_{work}$ | 0.193 | 0.142 |
| $R_{free}$ | 0.229 | 0.158 |
| No. of Atoms: | | |
| Protein | Chain A, 1163; Chain B, 1157 | 2989 |
| Water | 283 | 144 |
| Metal | 4 | 3 |
| Glycerol | --- | 28 |
| Average *B*-factor | 16.1 | 19.3 |
| **Validation** | | |
| RMSD bond lengths (Å) | 0.005 | 0.007 |
| RMSD bond angles (°) | 1.19 | 1.34 |
| Molprobity score (percentile) | $100^{th}$ | $98^{th}$ |
| Ramachandran favoured (%) | 99 | 98 |

*Data in parentheses is for the high-resolution shell.

**Supplementary Table 1. Data collection, refinement, and validation statistics for T450C and T450S X-ray crystal structures**

| Simulation type | System (mutant or wild type) | Number of replicates | Sampling per replicate (ns) |
|---|---|---|---|
| Molecular dynamics | D577H | 5 | 250 |
| | T450S/H572E | 5 | 250 |
| Metadynamics | Wild type | 3 | 6,000 |
| | T450S | 3 | 6,000 |

**Supplementary Table 2. Molecular dynamics and metadynamics replicate and sampling parameters.**

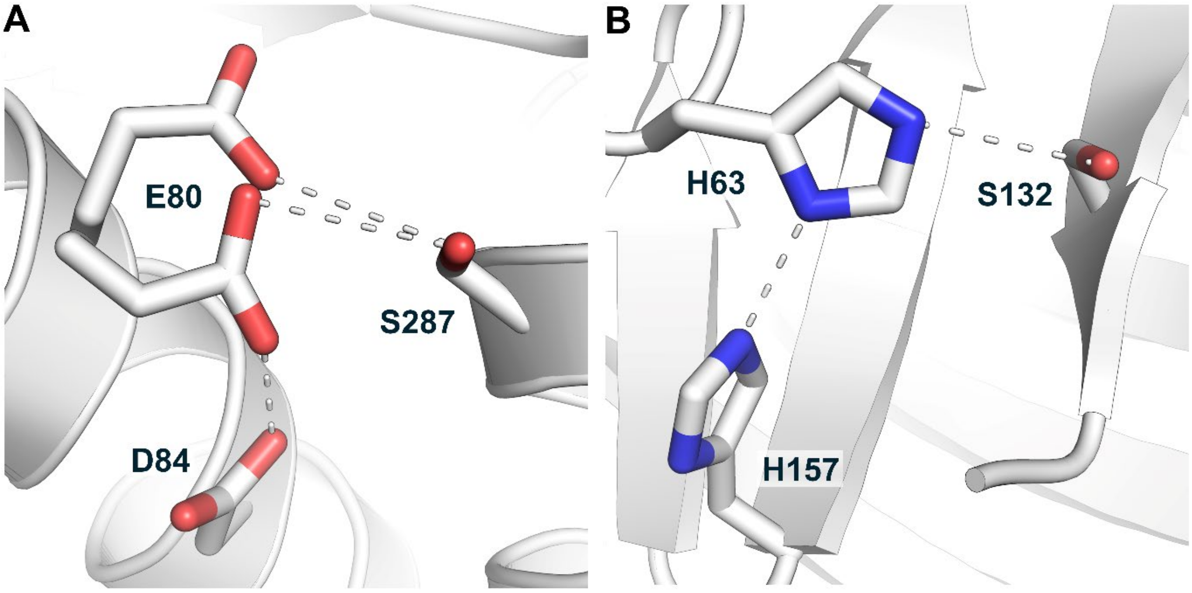


**Figure S1. The active site triads of two non-cannonical serine proteases. A.** Sedolisin, a serine protease from *Pseudomonas sp.* (PDB 1KDV), replaces the canonical histidine base of a serine protease with a glutamate (E80). **B.** The human beta herpesvirus 5 (PDB 1CMV) protease has a histidine, H157 in the classical acid position of the serine protease catalytic triad. This protease is also functional with a simpler dyad of H63/S132.

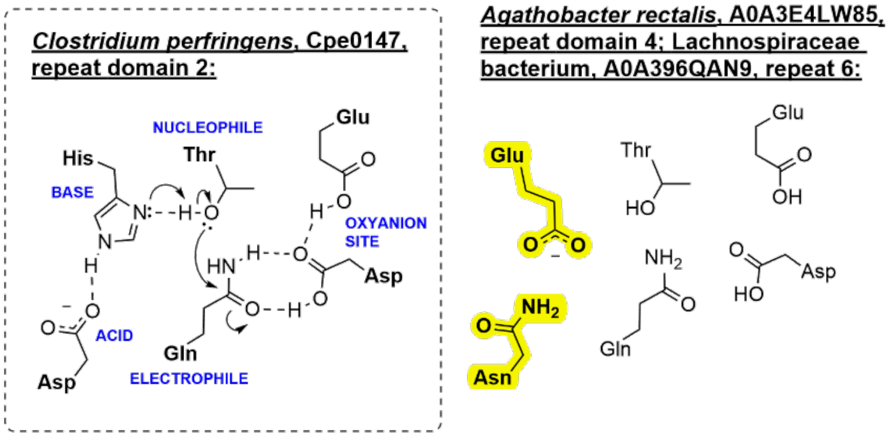


| Protein: | UNIPROT Code: | Role and structure: | Taxonomy:* | Structural Source: | Catalytic machinery: | |
|---|---|---|---|---|---|---|
| *Agathobacter rectalis*, A0A3E4LW85 repeat domain 4 [Pro596-Pro719] | A0A3E4LW85 | VaFE repeat-containing surface-anchored protein, mixed isopeptide & ester repeat domains (total 8) | Bacillota, Clostridia, Eubacteriales, Lachnospiraceae, *Agathobacter, A. rectalis* | AlphaFold | Acid<br>Base<br>Nucleophile<br>Electrophile<br>Oxyanion site | Asn<br>Glu<br>Thr<br>Gln<br>Asp, Glu |
| *Lachnospiraceae bacterium,* A0A396QAN9, repeat domain 6 [Pro730-Ile850]: | A0A396QAN9 | VaFE repeat-containing surface-anchored protein, mixed isopeptide & ester repeat domains (total 7) | Bacillota, Clostridia, Eubacteriales, Lachnospiraceae, unclassified Lachnospiraceae | AlphaFold | Acid<br>Base<br>Nucleophile<br>Electrophile<br>Oxyanion site | Asn<br>Glu<br>Thr<br>Gln<br>Asp, Glu |

* Phylum, Class, Order, Family, Genus, Species

**Figure S2. Naturally occurring variations in the putative catalytic residues of Cpe0147-like ester crosslink domains**. The identification of possible autocatalytic residues was achieved by visual inspection of X-ray crystal structures and AlphaFold predicted models overlaid with the Cpe0147 domain, PDB 4NI6. Residues that deviate from the canonical *Clostridium perfringens*, Cpe0147 adhesin domain 2 configuration, are highlighted in yellow.

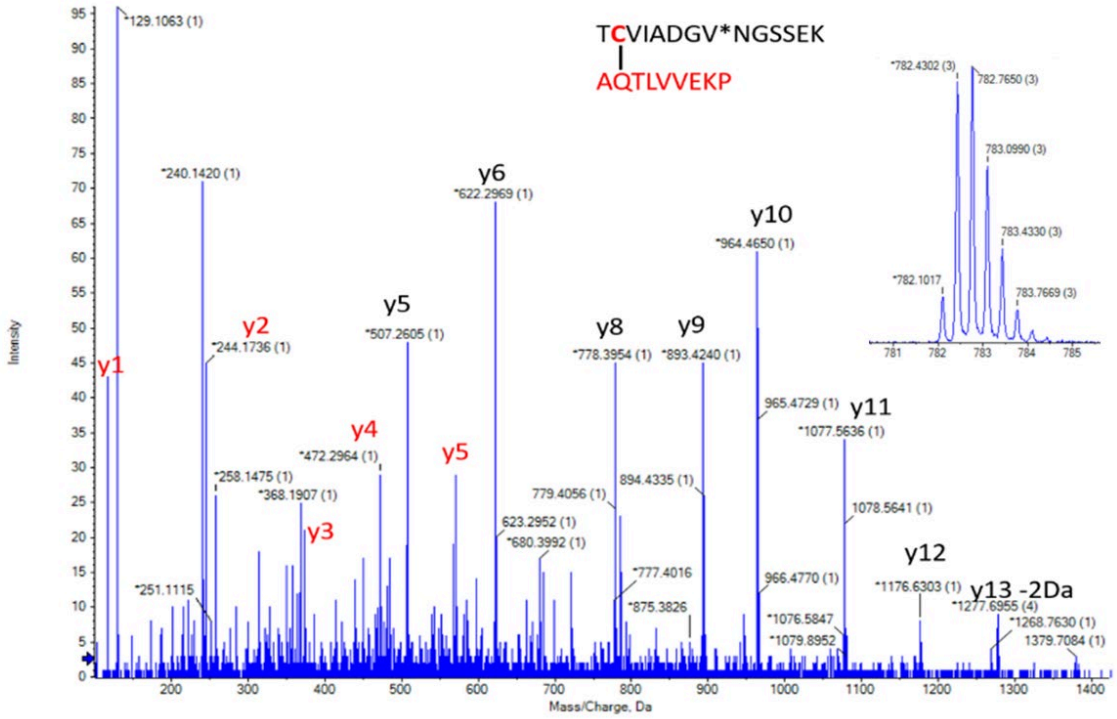


**Figure S3. Trypsin digest coupled with tandem mass spectrometry for T450C variant.** MS/MS spectrum of the m/z 782.77 3+ ion, representing the cross-linked peptides TCVIADGV*NGSSEK and AQTLVVEKP, with loss of ammonia (* indicates deamidated asparagine, +0.9848 Da). Charge states are indicated in parentheses. Inset shows the precursor ion spectrum.

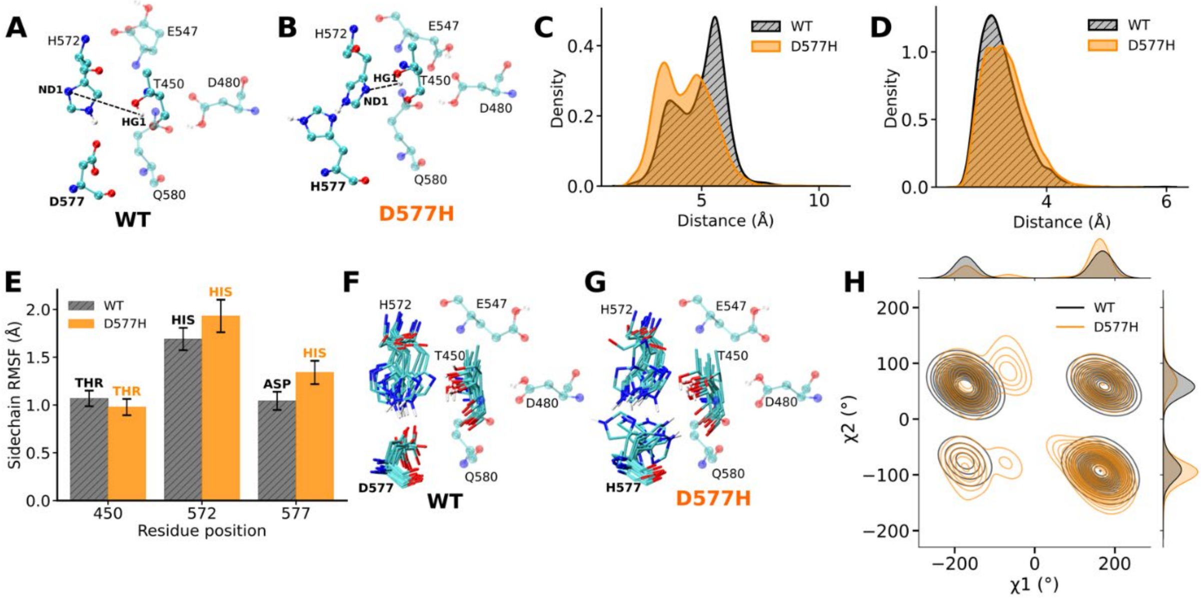


**Figure S4: Stabilising interactions for T-Q bond formation in Cpe0147. A-B** Representative conformations of WT (**A**) and D577H Cpe0147 mutant (**B**). The black dashed line indicates the distance between H572:ND1 and T450:HG1, suggested within the mechanism of bond formation. **C-D**. Probability density distributions of the H572:ND1—T450:HG1 and T450:OG1—Q580:NE2 distances, as sampled in MD simulations. **E**. Root mean square fluctuations (RMSF) of the residues suggested to influence the T-Q bond formation in the WT (grey) and D577H (gold) Cpe0147 constructs. **F-G**. Conformational ensembles of residues H572, D/H577 and T450, obtained from equilibrium molecular dynamics (MD) simulations of WT or D577H Cpe0147 constructs. **H.** Probability density function for H572 χ1 and χ2 dihedral angles, obtained from equilibrium simulations of WT (grey) and D577H (gold) Cpe0147 constructs.

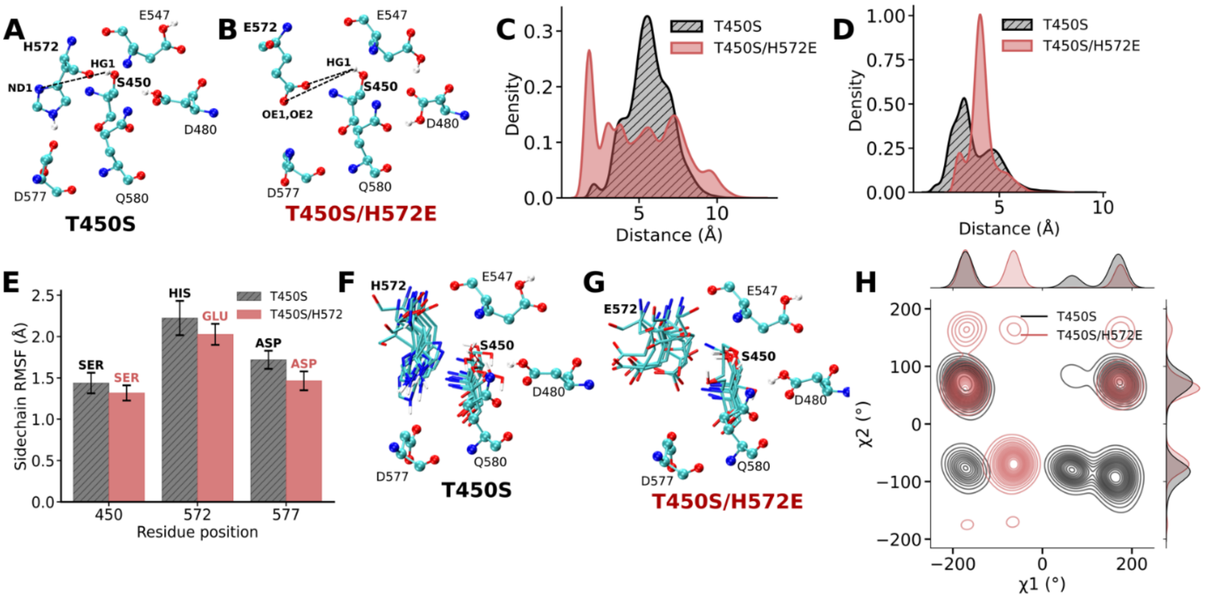


**Figure S5: Stabilising interactions for S-Q bond formation in Cpe0147. A-B.** Representative conformations of T450S (**A**) and T450S/H572E Cpe0147 mutants (**B**). The black dashed line indicates the distance between H572:ND1 and S450:HG1 (**A**), or E572:OE1,OE2 and S450:HG1 (**B**), suggested within the mechanism of bond formation. **C-D.** Probability density distributions of the H572:ND1—S450:HG1 and S450:OG1—Q580:NE2 distances, as sampled in MD simulations. **E.** Root mean square fluctuations (RMSF) of the residues suggested to influence the Ser-Gln bond formation in the T450S (grey) and H572E (pink) Cpe0147 constructs. **F-G.** Conformational ensembles of residues H/E572, D577 and S450, obtained from equilibrium molecular dynamics (MD) simulations of T450S or T450S/H572E Cpe0147 constructs. **H.** Probability density function for H572 and E572 $\chi 1$ and $\chi 2$ dihedral angles, obtained from equilibrium simulations of T450S (grey) and H572E (pink) Cpe0147 constructs.
